# Unraveling the Role of Urea Hydrolysis in Salt Stress Response during Seed Germination and Seedling Growth in *Arabidopsis thaliana*

**DOI:** 10.1101/2024.02.20.581165

**Authors:** Yuanyuan Bu, Xingye Dong, Rongrong Zhang, Xianglian Shen, Yan Liu, Shu Wang, Tetsuo Takano, Shenkui Liu

## Abstract

Urea is intensively utilized as a nitrogen fertilizer in agriculture, originating either from root uptake or from catabolism of arginine by arginase. Despite its extensive use, the underlying physiological mechanisms of urea, particularly its adverse effects on seed germination and seedling growth under salt stress remains unclear. In this study, we demonstrate that salt stress induces excessive hydrolysis of arginine-derived urea, leading to an increase in cytoplasmic pH within seed radical cells, which, in turn, triggers salt-induced inhibition of seed germination (SISG) and hampers seedling growth. Our findings challenge the long-held belief that ammonium accumulation and toxicity are the primary causes of SISG, offering a novel perspective on the mechanism underlying these processes. This study provides significant insights into the physiological impact of urea hydrolysis under salt stress, contributing to a better understanding of SISG.

## Introduction

Urea is the most widely used nitrogen fertilizer in agriculture globally (http://faostat.fao.org). About half of all nitrogen used for crop production is applied as urea. Urea is a primary nitrogen source taken up actively by plants from the soil solution. However, it is also an intermediate of plant arginine catabolism involved in nitrogen remobilization from source tissue. Despite this, the adverse effects of urea fertilizer on seed germination and seedling growth are still not reasonably explained. Therefore, the hydrolysis, transport, and utilization of plant urea, especially in extreme soils (saline-alkali soils), must be investigated better to develop strategies for knowledge-based crop improvement.

Arginase (ARGAH; arginine amidinohydrolyase), the only enzyme in plants known to generate urea *in vivo*, and plays a pivotal role in the mobilization of seed storage reserves during early seedling growth post-germination. The catabolism of arginine by arginase is crucial for mobilizing stored nitrogen upon germination and redistributing nitrogen from source tissues, as evidenced by various studies (Goldraij et al., 1999; Todd et al., 2001; Todd et al., 2002). Arginine, abundant in the storage proteins of most plant seeds, has the highest nitrogen-to-carbon ratio (N:C = 4:6) among all amino acids, making it a significant nitrogen source for seed reserve mobilization (Jones et al., 1968; King et al., 1997; Polacco et al., 2013). In a study of 379 angiosperm seeds, arginine accounted for 17% of the seed nitrogen content (8.58 g arginine/16 g seed N) (Van Etten et al., 1967).

Beyond its hydrolysis by arginase to yield urea, arginine also serves as a substrate for nitric oxide (NO) and polyamine (PA) biosynthesis in plants by nitric oxide synthase (NOS) and arginine decarboxylase (ADC) (Siddappa et al., 2020), respectively. Consequently, arginine becomes a shared substrate, competitively utilized by the three enzymes ARGAH, NOS and ADC (Fig. 1). Many recent studies suggested that the improved salt tolerance observed in arginase-deficient mutants is attributed to an indirect upregulation of NOS and ADC pathways, thereby enhancing NO and PA-mediated plant defense responses (Flores et al., 2008; Shi et al., 2013; She et al., 2017; Wang et al., 2011a; Winter et al., 2015). However, this raises the question of how the presence of arginase activity still allows to urea-deficient mutants to significantly alleviate salt-induced inhibition of seed germination (SISG) and bolster seedling growth (Bu et al., 2015).

**Figure 1.**
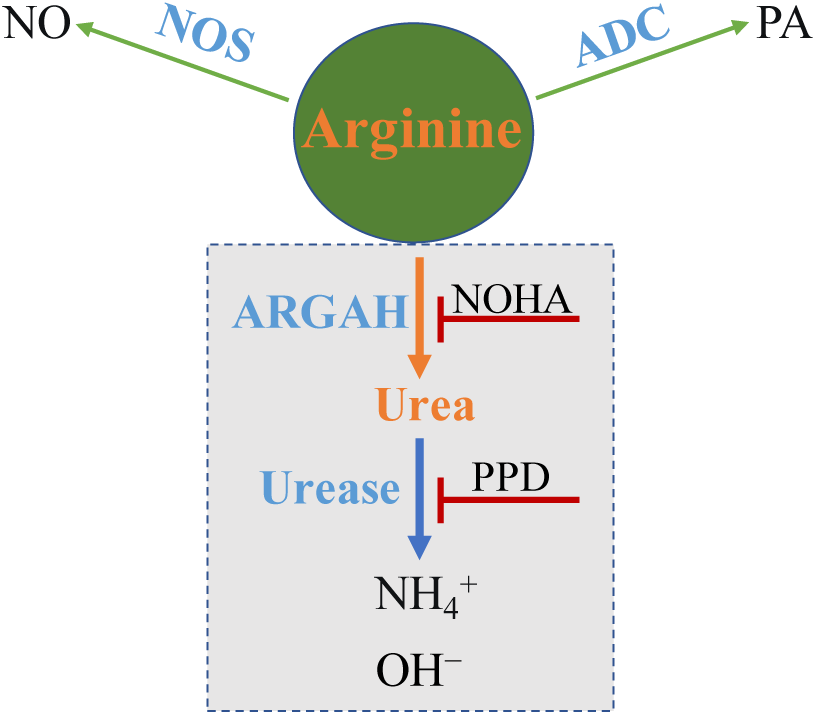
A simple model of arginine metabolism in *Arabidopsis thaliana*. This model outlines the conversion of arginine into: (1) nitric oxide (NO) and citrulline by nitric oxide synthase (NOS); (2) polyamine (PA) by arginine decarboxylase (ADC); and (3) ornithine and urea by arginase, with urea further decomposed to ammonia by Urease. It highlights arginine as a shared competitive substrate for the three enzymes ARGAH, NOS and ADC, illustrating the competitive enzymatic interactions. NOHA: *N*^G^-hydroxy-L-arginine, an arginase inhibitor; PPD: phenyl phosphorodiamidate, a Urease inhibitor.

To address above these questions, our study focused on the connection between urea produced by arginase hydrolysis pathway, distinct from the PA and NO pathways, and the salt-induced inhibition of seed germination and seedling growth, with a particular focus on the dual-step hydrolysis of arginine (Fig. 1) and the internal transport of urea in relation to SISG and seedling growth. By employing specific enzyme inhibitors and gene deletion mutants within the arginine two-step hydrolysis pathway, we found that hydrolysis of arginine-derived urea is the key point for effectively alleviating the adverse effects of salt on seed germination and seedling growth. This led us to question: what mechanism underlies the triggering of SISG and stunted seedling growth by urea hydrolysis? We ruled out the potential accumulation of ammonium, resulting from urea hydrolysis, by incorporating exogenous ammonium into the salt stress medium. Interestingly, our findings on intracellular pH measurements indicate that the salt-induced hydrolysis of urea, yielding OH^-^, leads to an increase in intracellular pH of the radicle, pinpointing this as the primary factor initiating SISG and impeding seedling growth. These insights provide a novel understanding of mechanisms behind SISG and seedling development.

## Results

### Urea hydrolysis is the cause of SISG in the two-step arginine hydrolysis

Initially, to determine whether arginase-mediated arginine hydrolysis pathway is involved in SISG and seedling growth, we utilized two inhibitors specific to the two-step hydrolysis reaction of the arginine hydrolysis pathway: *N*^G^-Hydroxy-L-arginine (NOHA), an arginase inhibitor, and Phenyl phosphorodiamidate (PPD), a urease inhibitor (Fig. 1). We analyzed the seed germination phenotypes under these conditions. In the absence of NOHA, the NaCl treatment significantly hindered the seeds’ capacity to germinate, as evidenced by the radicle’s inability to to break through the seed coat (Fig. 2a). However, the addition of 5 μM NOHA into NaCl medium significantly increased the germination rate (Fig. 2a). In contrast, the presence or absence of NOHA in the control medium did not notably affect seed germination (Fig. 2a).

**Figure 2.**
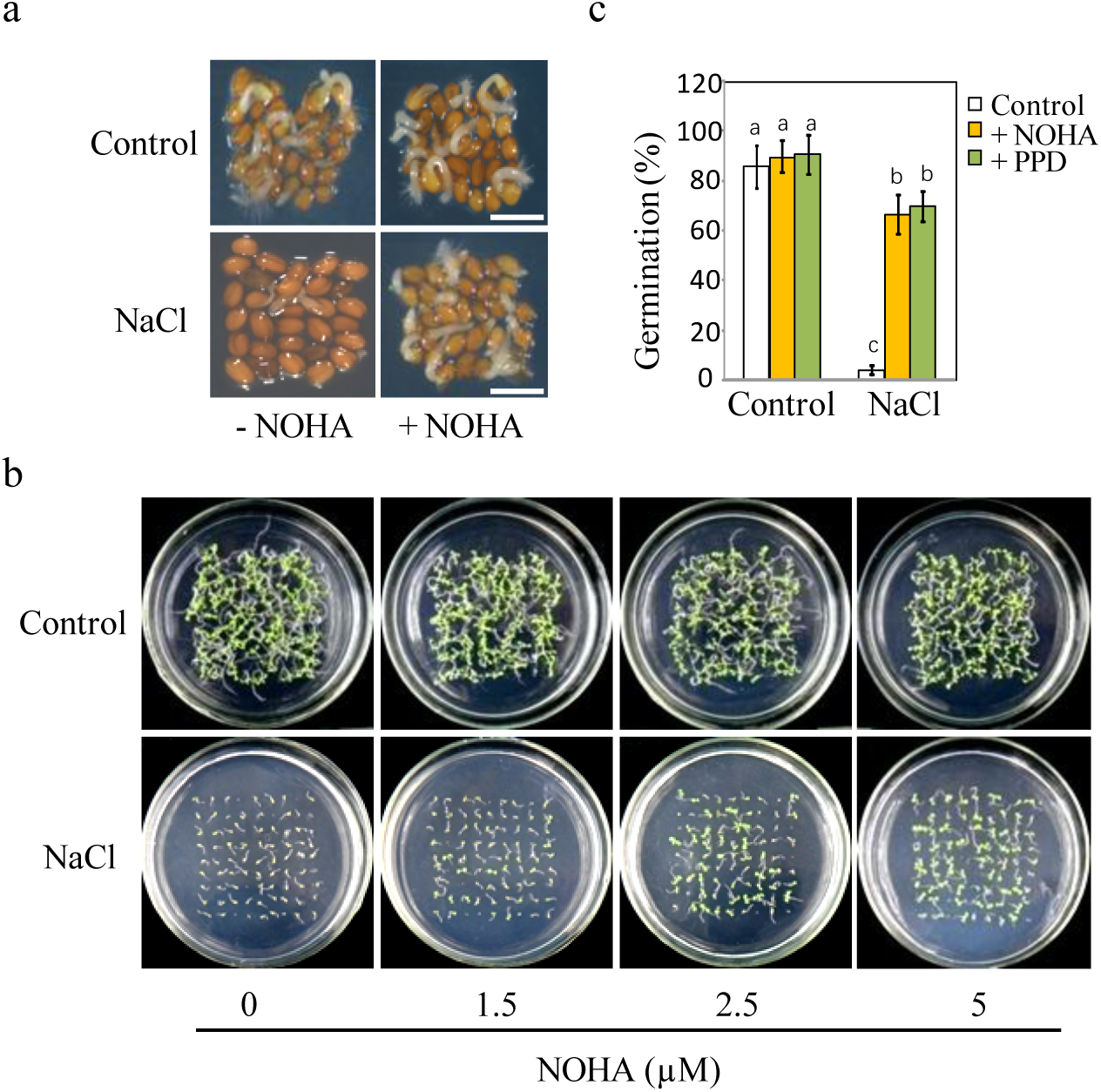
Blocking arginine hydrolysis and promoting seed germination under salt stress. (a) Comparison of germination rates of WT seeds on half-strength MS (½ MS) medium containing 0 mM (Control) or 135 mM NaCl with 5 μM NOHA or without NOHA. Photographs were taken at the 48 h after stratification. A representative result from one of three independent experiments, all yielding similar outcomes, is shown. Scale bar represents 1 mm. (b) Seeds were germinated on ½ MS medium containing either 0 mM or 135 mM NaCl and varying concentrations of NOHA (0, 1.5, 2.5, 5 μM); photographs were taken 14 days post-germination. (c) Germination rates of WT seeds in either 0 mM or 135 mM NaCl medium with or without 5 μM NOHA or 15 μM PPD. Stratification consisted of pretreatment of seeds for 2 days at 4 °C in the darkness. In all experiments, seeds were freshly sowed and incubated under 16-h light and 8-h dark conditions at 22°C. The experiment was repeated three times; at least 30 seeds were counted in each replicate. The data was analyzed using a one-way analysis of variance (ANOVA), followed by Duncan’s multiple range test for the post-hoc comparisons. Significant differences between groups, indicated by different letters on the error bars, were determined (*P*< 0.05).

To better observe the effect of NOHA on seed germination or early seedling growth, different concentrations of NOHA (0, 1.5, 2.5, and 5µM) were added to the control or NaCl mediums (Fig.2b). The results showed that the inhibitory effect of salt on seed germination was significantly alleviated by the increasing NOHA concentrations, while NOHA added in control medium had no substantial effect (Fig.2b). Additionally, root length measurements taken 14 post-germination indicated that NOHA could promote root growth under salt stress conditions (supplementary Fig.1).

To directly elucidate the impact of the arginase-mediated arginine hydrolysis pathway on SISG, and under the assumption that the competition for arginine substrate by multiple pathways (Fig. 1) is not affected at this stage, we investigated the effect of urea hydrolase inhibitor PPD, which acts on a downstream metabolite of arginine, on seed germination. This was then compared to the effects observed in the NOHA assay. The results showed that salt significantly inhibited the germination of wild-type Arabidopsis seeds, with less than 10% germination. The addition of 15 µM of PPD increased the germination rate to 70%, compared with the rate without PPD (Fig. 2c), which showed the same trend as NOHA treatment (Fig. 2c). These results indicate that salt tolerance in WT seeds germination and seedling growth is manifested especially under NOHA and PPD conditions. The two-step hydrolysis pathway of arginine mediated by arginase and its downstream metabolite urea hydrolase is not only involved in SISG events, but also urea hydrolysis may be the leading cause of SISG occurrence.

### Genetic evidence for SISG triggered by the arginine hydrolysis pathway

To clarify the genetic basis of arginase inhibitor experiments described above, the following studies were conducted. *Arabidopsis thaliana* has two genes encoding ARGAH, namely ARGAH1 (gene number: AT4G08900) and ARGAH2 (gene number: AT4G08870). *AtArgAH1* has a T-DNA insertion within its first exon, while *AtArgAH2* contains a T-DNA insertion in the fourth exon. Consequently, the mutants resulting from these insertions were denoted as *argah1* and *argah2*, respectively (Supplemental Fig. 2a). Additionally, *AtArgAH1* and *AtArgAH2* double knockout mutants (*argah1/argah2*) were gnerated using clustered regularly interspaced short palindromic repeats (CRISPR)/of CRISPR-associated (Cas) protein 9 system. Targeted modifications were introduced into the third exon of the *AtArgAH1* gene and the second exon of the *AtArgAH2* gene. In total, four Hygromycin-resistant lines were obtained from the F_0_ transgenic lines. Subsequent to CRISPR-Cas9 gene editing and sequencing, one line of the *argah1/argah2* double mutant exhibited a 1-bp insertions in the coding region of the *AtArgAH1* gene, and another 1-bp insertions in the coding region of the *AtArgAH2* gene (Supplemental Fig. 2b). Thus, for subsequent genetic phenotypic analyses, we used *argah1/argah2* double mutants were utilized alongside wild-type Arabidopsis (WT). Seed germination and initial growth post-germination of all mutants (*argah1, argah2*, and *argah1/argah2*) were not significantly different from those of WT in the control medium (Fig. 3a). In contrast, adding NaCl to the control medium severely inhibited seed germination and radicle development of WT (Fig. 3b). The seed germination rates of each mutant (*argah1, argah2*, and *argah1/argah2*) were significantly higher than those of WT under NaCl conditions (Fig. 3c). Furthermore, the root length of the mutant on day 14 post-seed germination was significantly higher than that of the WT (Fig. 3d). Moreover, arginase activity analysis revealed significantly lower levels in *argah1, argah2*, and *argah1/argah2* mutants following NaCl treatment compared to WT (Fig. 3e). These findings indicate that partial or complete blocking of the arginine hydrolysis pathway can effectively alleviate SISG. We have previously reported that deletion mutants of the urea hydrolase gene (*urease*) enhances salt tolerance during seed germination (Bu et al., 2015). That implies that even the accumulation of urea *in vivo*, derived from the hydrolysis of arginine by arginase, is insufficient to trigger SISG.

**Figure 3.**
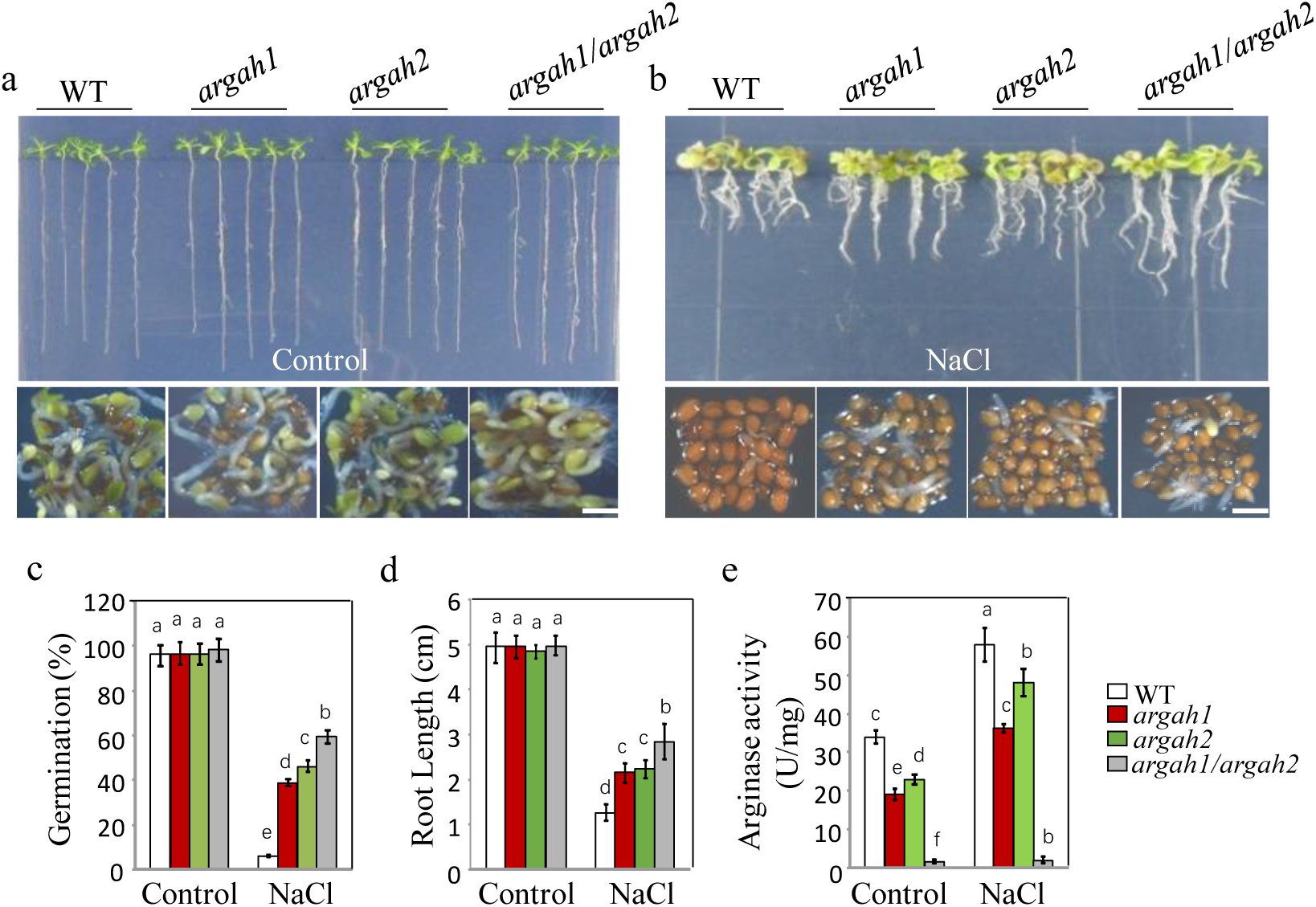
Deletion of arginase hydrolytic pathway alleviates the inhibition of seed germination by salt. WT, *argah1*, *argah2* and *argah1/argah2* mutant seeds germinated, and seedling growth was observed on ½ MS medium under two conditions: 0 mM NaCl (control) (a) and 135 mM NaCl (b). Photographs captured the seedlings 14 days post-germination (top) and 48 hours post-stratification (bottom). (c) germination rates, (d) root length measurements, and (e) arginase activity assays for WT, *argah1*, *argah2*, and *argah1/argah2* mutants on control and 135 mM NaCl mediums. In all experiments, seeds were freshly sowed and incubated under 16-h light and 8-h dark conditions at 22°C. The experiment was repeated three times; at least 30 seeds were counted in each replicate. Scale bar: 1 mm. Data were subjected to a one-way analysis of variance, followed by Duncan’s post-hoc test. Different letters on the error bars denote significant differences in the data (*P*< 0.05).

It is well known that Na^+^ toxicity is the main factors triggering SISG. The mechanism involves SOS3 binds to free Ca^2+^, consequently activating SOS2 protein kinase, which in turn phosphorylates SOS1, leading to the activation of SOS1 transport responsible for pumping Na^+^ out of the cell. Therefore, SOS3-deficient mutant (*sos3*) accumulate substantial Na^+^ levels and are highly sensitive to salt stress. In order to further clarify the importance of urea hydrolysis in SISG, *sos3* mutants were considered as standard materials for SISG analysis, along with *aturease/sos3* double mutant. Initially, we introduced PPD to 50 mM NaCl medium to observe the growth of *sos3*. Notably, with increasing the concentrations of PPD, the growth inhibition of *sos3* induced by NaCl was alleviated significantly (Fig. 4a). Furthermore, both seed germination rate (Fig. 4b) and 14-d root length (Fig. 4c) were significantly increased compared to those without PPD treatment. Subsequently, the *aturease/sos3* double mutant analysis revealed that significant improvement in germination ability and growth compared to *sos3* in the presence of 50 mM NaCl (Fig. 4d and e). In addition, the root length of *aturease/sos3* double mutant exceeded that of *sos3* mutants under salt stress conditions (Fig. 4f). The above treatments with urea hydrolase inhibitors PPD, coupled with genetic evidence, suggest that blocking the hydrolysis of arginine-derived urea can mitigate the hypersensitivity of *sos3* to salt stress. This further corroborates that urea accumulation *in vivo* is not the primary trigger of SISG. Instead, urea hydrolysis appears to play a dominant role in this process.

**Figure 4.**
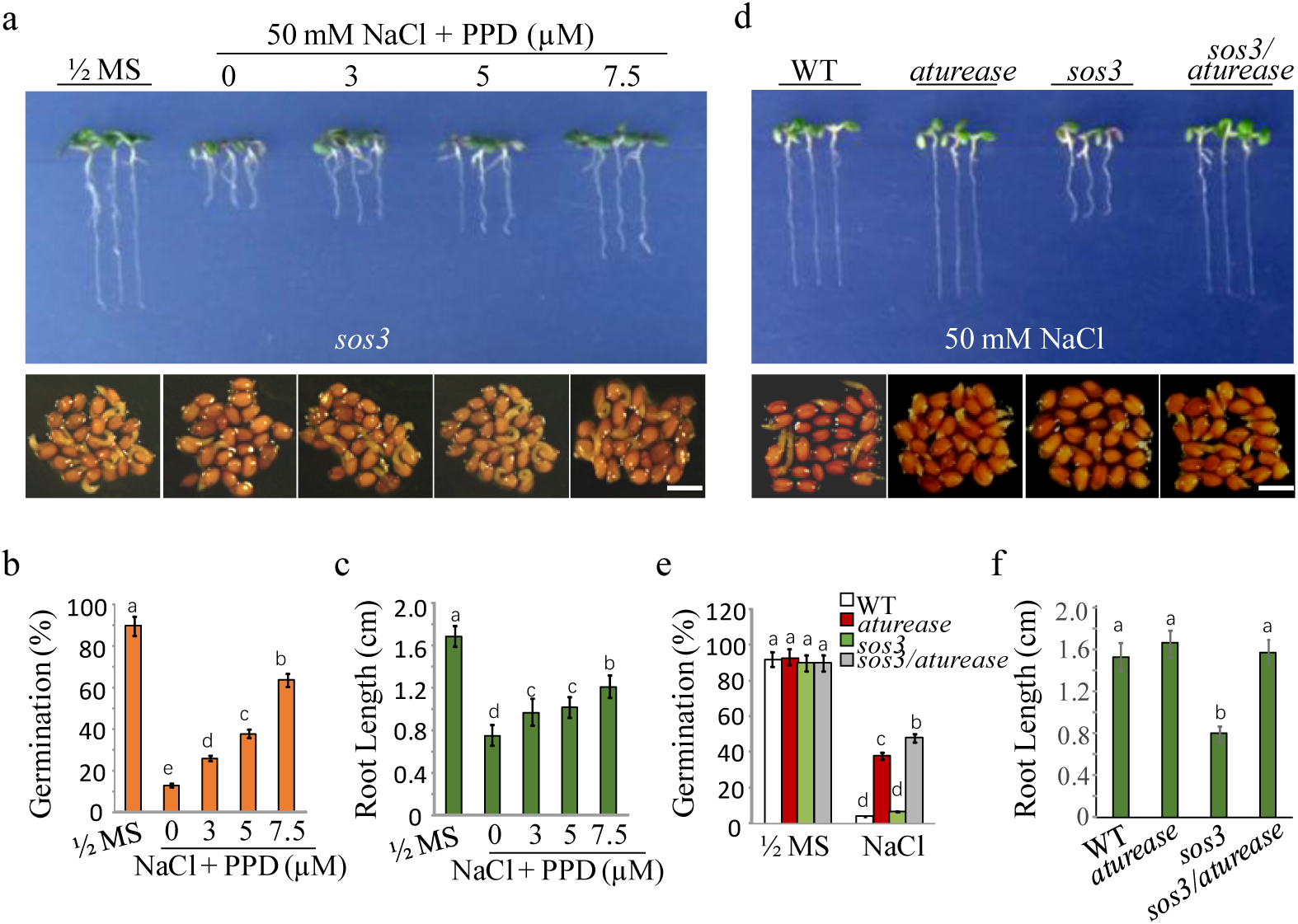
Blocking the arginine hydrolysis pathway mitigates the hypersensitivity of *sos3* mutants to salt. (a) seeds germinated and seedling growth of *sos3* mutants on ½ MS with 50 mM NaCl medium, supplemented with PPD at different concentrations (0, 3, 5, 7.5 μM). Images captured at 14 days post-germination (top) and 48 hours post-stratification (bottom). (b) Germination rates and (c) root lengths of *sos3* mutants assessed after the specified treatments. (d) WT, *aturease*, *sos3* and *sos3/aturease* seeds germinated and grew on ½ MS with 50 mM NaCl. Photographs were taken 14 days post-germination (top) and 48 hours post-stratification (bottom). (e) Germination rate and (f) root lengths for WT, *sos3*, *aturease* and *sos3/aturease* mutants were measured after the indicated treatments. In all experiments, seeds were freshly sowed and incubated under a 16-hour light/8-hour dark cycle at 22°C. The experiment was repeated three times; at least 25 seeds were counted in each replicate. Scale bar: 1 mm. The data were analyzed using one-way ANOVA followed by Duncan’s post-hoc test, with different letters on the error bars indicating significant differences in the data (*P*< 0.05).

### How does the hydrolysis of arginine-derived urea trigger SISG

Next, if urea accumulation is not the key to SISG, is urea hydrolysis detrimental to seed germination? The cause of SISG by excessive urea hydrolysis in triggering SISG remains an enigmatic issue. Initially, we confirmed that NaCl stress resulted in the accumulation of urea (Fig. 5a) and NH_4_ ^+^ (Fig. 5b) during WT seed germination, suggesting that salt stress promoted seed nitrogen mobilization and significantly lowered urea and NH_4_ ^+^ levels in the *argah1/argah2* double mutants (Fig. 5a and b). In addition, in the presence of 50 mM NaCl, *sos3*-mutant seeds accumulated higher NH_4_ ^+^ levels than WT and *aturease* mutants. Remarkably, the *aturease/sos3* double mutant effectively reduced NH_4_ ^+^ levels to those akin to WT and *aturease* (Fig. 5c). These findings initially indicated that SISG might be triggered by the ammonium accumulation resulting from excessive hydrolysis of arginine-derived urea. However, the following experiments challenged this seemingly reasonable hypothesis. We examined seed germination and growth in the presence of both NaCl and NH_4_Cl. Surprisingly, exogenous NH_4_ ^+^ significantly alleviated the inhibitory effects of NaCl on seed germination (Fig. 5d and Supplementary Fig. 3), with a concomitant increase in NH_4_ ^+^ (Fig. 5e), indicating that NH_4_ ^+^ accumulation was not the primary cause of SISG. Further validation was sought by adding urea to NaCl medium. Intriguingly, urea substantially inhibited seed germination (Fig. 5d and Supplementary Fig. 3), but NH_4_ ^+^ content did not accumulate in seeds (Fig. 5e), thereby confirming that NH_4_ ^+^ produced via urea hydrolysis was not the driving force behind SISG.

**Figure 5.**
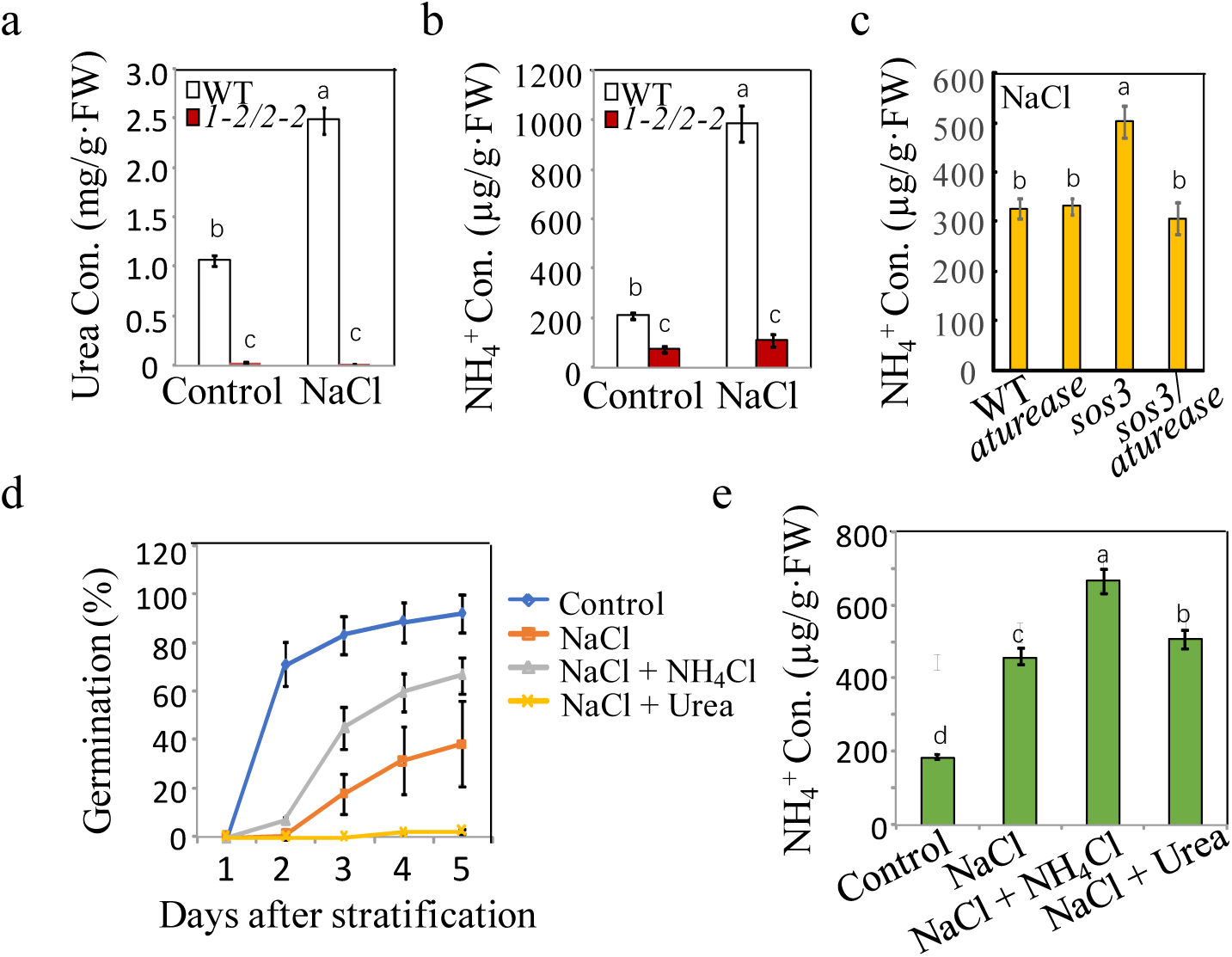
Correlation between metabolites from arginine hydrolysis and salt inhibit-induced inhibition of seed germination. (a) Urea and (b) NH ^+^ concentrations were measured in WT and *argah1/argah2* seedlings grown in ½ MS medium under Control (0 mM) and 135 mM NaCl conditions. (c) NH ^+^ levels assessed in WT, *aturease*, *sos3* and *sos3/aturease* seedlings under 50 mM NaCl treatment in ½ MS containing. (d) Effect of NH_4_Cl on salt response was evaluated in WT seeds, monitoring germination rates in control medium with 135 mM NaCl, with and without 10 mM NH_4_Cl and 10 mM Urea, over time. (e) NH ^+^ concentration in WT seedlings measured on ½ MS with 135 mM NaCl, with or without the addition of 10 mM NH_4_Cl and 10 mM Urea. In all experiments, seeds were freshly sowed and incubated under a 16-hour light/8-hour dark cycle at 22°C. The experiment was repeated three times; at least 30 seeds were counted in each replicate. Scale bar: 1 mm. The data were analyzed using a one-way ANOVA, with Duncan’s test for post-hoc comparison. Different letters on the error bars indicate significant differences in the data (*P*< 0.05).

These results disproved that NH_4_^+^ accumulation and toxicity caused by urea hydrolysis was the primary cause of SISG. Then, we reconsidered the hydrolysis process of urea, based on the complete reaction of urea hydrolysis: (NH_2_)_2_CO + 3H_2_O → 2NH_4_ ^+^ + HCO_3_^-^ + OH^-^, and discovered that in addition to NH_4_ ^+^, excessive urea hydrolysis by alkaline reaction producing a considerable accumulation of OH^-^. According to recent reports, excessive assimilation of NH_4_ ^+^ by glutamine synthetase (GS) produces acid stress and has toxic effects on plants (Hachiya et al., 2021; Witte et al., 2011). To verify the relationship between acid stress of NH_4_ ^+^ assimilation by GS and plant salt sensitivity, we added GS inhibitor MSX to a salt-stress medium to inhibit NH_4_ ^+^ assimilation (Kiyomiya et al., 2001; Rawat et al., 1999) and found that MSX significantly inhibited the germination of WT seeds under salt stress, while WT germinated well in the control medium (Fig. 6a and b). In addition, the percentage of seedlings with green cotyledons (Fig. 6c) and root growth (Fig. 6d) were severely restricted under salt-stress conditions. These results suggest that the acidification process of GS assimilation is beneficial to plant resistance to salt stress.

**Figure 6.**
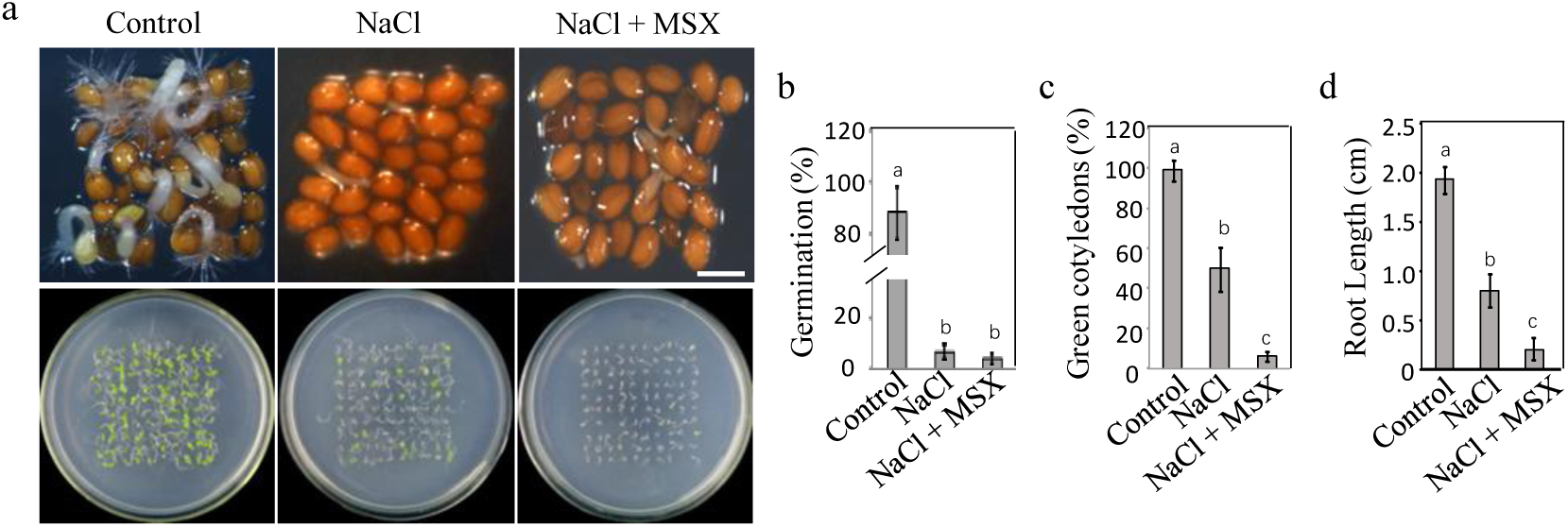
Effect of acidification process of glutamine synthetase assimilation on plant salt tolerance. (a) germination of WT seeds on ½ MS medium (Control), ½ MS medium supplemented with 135 mM NaCl, and ½ MS medium containing 3 µM MSX, with photographs taken 48 h post-stratification (bottom) and 10 days post-germination (top), scale bar representing 1 mm. The study evaluated (b) WT seeds germination rates, (c) the count of green cotyledons, and (d) root length. In all experiments, seeds were freshly sowed and incubated under 16-hour light and 8-hour dark conditions at 22°C. The experiment was repeated in triplicate with a minimal 30 seeds per replicate. The data analysis was performed using a one-way ANOVA and Duncan’s post-hoc test, with significant differences (*P*< 0.05) denoted by different letters on the error bars.

If the acidification process promotes plant resistance to salt stress, it follows that alkaline may exacerbate the inhibition of salt stress on seed germination and growth. Given that urea hydrolysis is an alkaline reaction process, we further conducted a thorough analysis of its impact on intracellular pH under salt stress. Initially, Arabidopsis seeds expressing a fluorescent pH indicator (PRpHluorin) were treated with NaCl and PPD (see Materials and Methods). To establish a correlation between PRpHluorin fluorescence ratios and pH values, an *in vivo* calibration was performed across the pH range indicated in Figure 7a. We reasoned that *in vivo* calibration ensured that obtained pH ratios accurately reflected the intracellular environment, facilitating robust measurements of cytoplasmic pH over the pH range of 6 to 8. Then, we qualitatively observed panoramic pH in germinating seeds revealed a trend towards alkalinization in NaCl-treated seeds compared to the control (no salt treatment), especially evident in the epidermal cells of the root elongation zone (Fig. 7b). However, the addition of PPD effectively alleviated this alkalinization of the cells under salt treatment conditions, restoring their pH to levels similar to the control.

**Figure 7.**
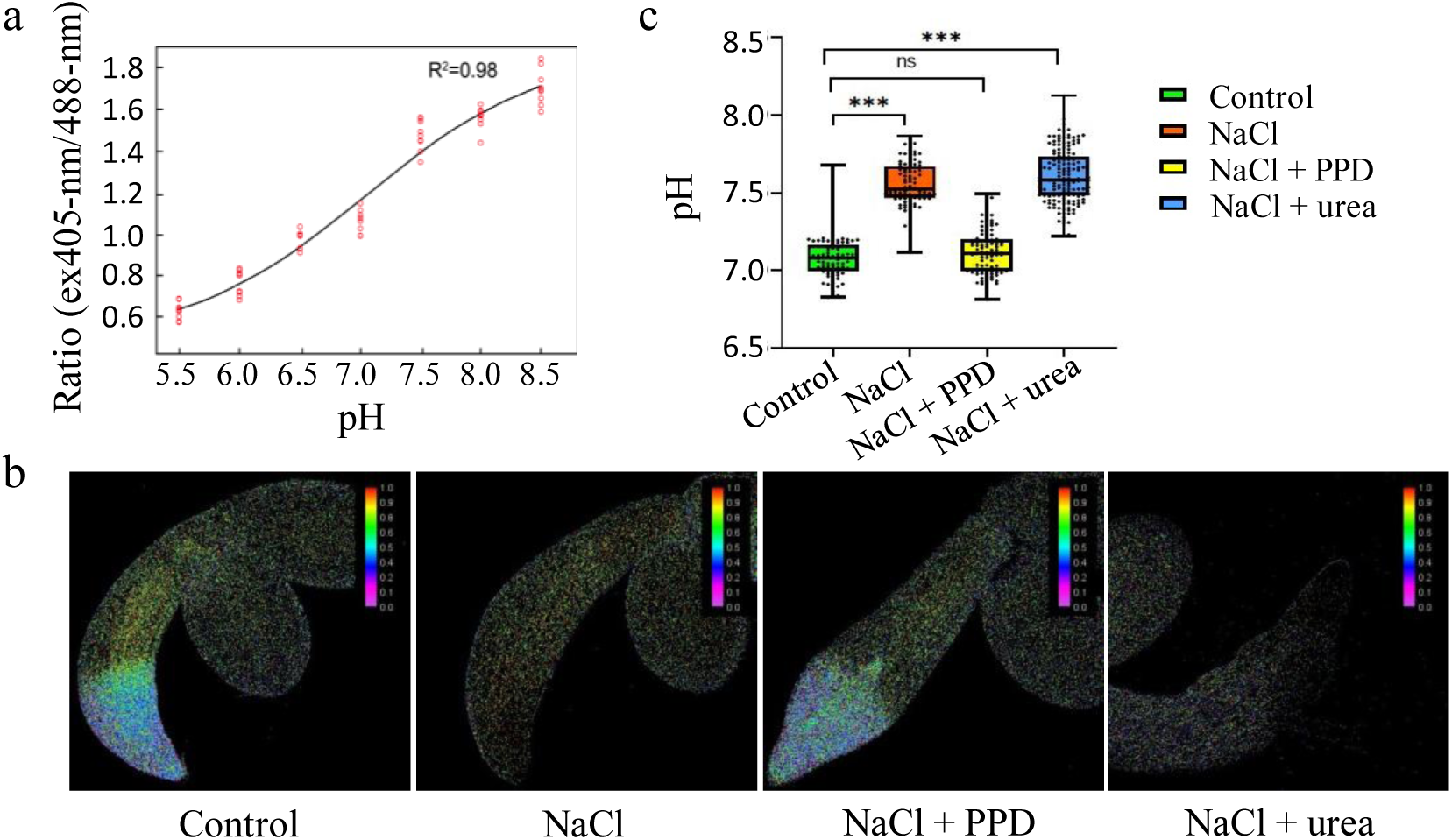
Inhibition of urea *via* arginine hydrolysis promotes seed germination under salt stress by lowering cell pH. (a) calibration curve for pHluorin in seedlings adjusted to various pH levels using 50 mM Mes-BTP (pH 5.2–6.4), 50 mM Hepes-BTP (pH 6.8–7.6), and 50 mM ammonium acetate. The curve plots the average fluorescence intensity ratios against the pH for 15 seedlings. (b) fluorescence ratio images (emission 500 to 550 nm) of PRpHluorin expression seedlings grown in control, 135 mM NaCl medium, 135 mM NaCl with 15 µM PPD, and 135 mM NaCl with 10 mM urea, at 22°C for 3 days. Scale bar represents 100 μm. (c) Boxplots depict the cytoplasmic pH in root epidermal cells of seedlings, measured using the PRpHluorin fluorescence in the root elongation zone under treatments. Data points indicate individual pH measurements. Statistical significance was assessed by one-way ANOVA (* *P* <0.05, ** *P* < 0.01, *** *P* < 0.0001).

Subsequently, we quantified the pH of epidermal cells in the root elongation zone at the onset of seed germination (3 days old) for PRpHluorin seeds. Each group with no less than 10 samples was calculated to determine the intracellular pH of multiple cells in the elongated root epidermal region of germinated seeds. The results showed that the average cytoplasmic pH of the control was 7.07±0.02, while the salt treatment increased the pH to7.76±0.03. Conversely, salt treatment with PPD restored the pH to 6.97± 0.03 (Fig. 7c), akin to the control (Fig. 7b). Additionally, exogenous urea treatment led to a clear trend towards higher pH in root cells compared to the salt-only control (Fig. 7b and c), further affirming that the increase in intracellular pH induced by urea hydrolysis inhibited seed germination. These results validated our initial hypothesis that blocking urea hydrolysis by PPD reduces intracellular pH, effectively alleviating SISG. Excessive hydrolysis of arginine-derived urea elevates the cytoplasmic pH of seed radical cells, triggering SISG and impeding seedling growth in *Arabidopsis thaliana*.

### Blocking the transport of arginine-derived urea is beneficial for alleviating SISG

Based on our findings, we first summarized a hypothetical model of the arginine hydrolysis pathway that regulates seed germination under salt stress (Fig. 8). Salt stress highly induced the accumulation of arginine through the degradation of seed-stored proteins in the cotyledon. Then, arginine is hydrolyzed to urea through the action of arginase, and urea is further degraded by plant urease to form NH_4_ ^+^ and OH^-^ (Fig. 8a). During this process, the inhibition of salt stress on seed germination and seedling growth can be alleviated by blocking either of the two steps of the arginine hydrolysis pathway (Fig. 8b). The actual trigger for SISG is the increase in intracellular pH caused by the hydrolysis of urea produced by arginine in the root rather than the effect of NH_4_ ^+^. It is noteworthy that urea production in the cotyledon is transported to the root by urea transporter; however, urea transport is also worth exploring if the SISG events caused by the arginine hydrolysis pathway of seed storage nitrogen mobilization can be understood at the tissue and organ level.

**Figure 8.**
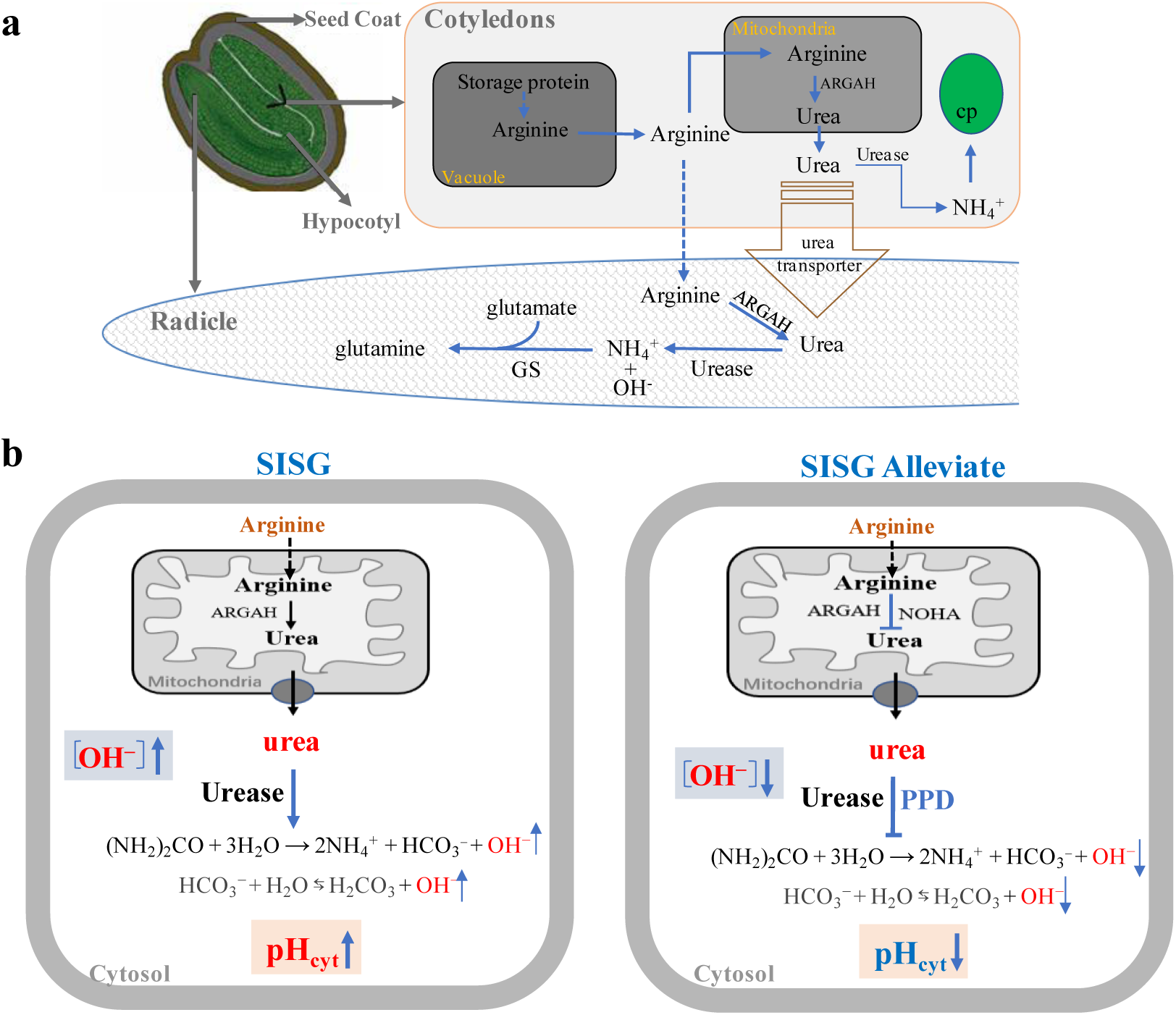
Hypothetical model for the regulation of seed germination by the Arginine hydrolysis pathway under salt stress condition. (a) Salt stress highly induced the accumulation of arginine through the degradation of seed-stored proteins in the cotyledon. Then, arginine is hydrolyzed to urea through the action of arginine amidinohydrolyase (ARGAH), and urea is further degraded by plant Urease to form NH_4_^+^ and OH^-^. (b) Salt stress stimulates the arginine hydrolysis pathway, leading to urea production by ARGAH, and its subsequent breakdown by Urease, increasing NH_4_^+^, HCO ^-^, OH^-^ levels, and cytoplasmic pH (pH), which inhibits seed germination (SISG). Application of arginine hydrolysis inhibitors (e.g., NOHA for arginase, and PPD for Urease) or using *ARGAH* or *Urease* gene deletion mutants under salt stress can reduce NH ^+^, HCO ^-^, OH^-^ levels, and decreased pH, thus promoting seed germination (SISG alleviate). The model delineates the urea hydrolysis process resulting in increased cytoplasmic pH, NH ^+^ and alkalinity, contributing to the inhibition of seed germination under salt conditions, abbreviated as SISG (salt inhibits seed germination). PPD, an inhibitor of urea hydrolase.

Here, we identified a loss-of-function mutant, *atdur3*, which lacks the urea transporter gene AtDur3 (Supplemental Fig. 4a∼c). The *atdur3* mutant line exhibited impaired growth on a medium containing urea as the sole nitrogen source (<5 mM) (data not shown). When *atdur3* and WT were germinated on control mediums, there was no significant difference in their seed germination ability and root development (Fig. 9a∼c). On the control medium supplemented with NaCl, the germination ability of WT seeds was significantly inhibited (the ability of radicle to break through the seed coat), and the germination rate of *atdur3* mutant seeds was significantly higher than that of WT (Fig. 9a and b). These results suggest that blocking urea transport *in vivo* could effectively alleviate SISG. We hypothesized that blocking the transport of urea from arginine hydrolysis in the cotyledon to the root and blocking its hydrolysis in the root would promote the germination of seeds under salt stress. Therefore, we subsequently analyzed urea concentrations in the roots and cotyledon of *atdur3* under salt stress. Under non-stressed normal conditions (control), root urea concentrations were lower in *atdur3* plants than in WT plants, but urea levels in cotyledons had no significant difference (Fig. 9d). However, when plants were grown under NaCl stress, the urea levels were lower in *atdur3* roots than in WT roots. Still, urea levels were higher in *atdur3* cotyledons than in WT cotyledons (Fig. 9d). We further analyzed the change ratio of urea (cotyledon/root), the results show that urea was mainly accumulated in atdur3 cotyledons under control or NaCl conditions (Fig. 9e). These results further indicate that the transport of urea, the primary metabolite of arginine hydrolysis, plays an important role in SISG, that is, blocking the transport of arginine-derived urea from the cotyledon to the root is conducive to the alleviation of SISG.

**Figure 9.**
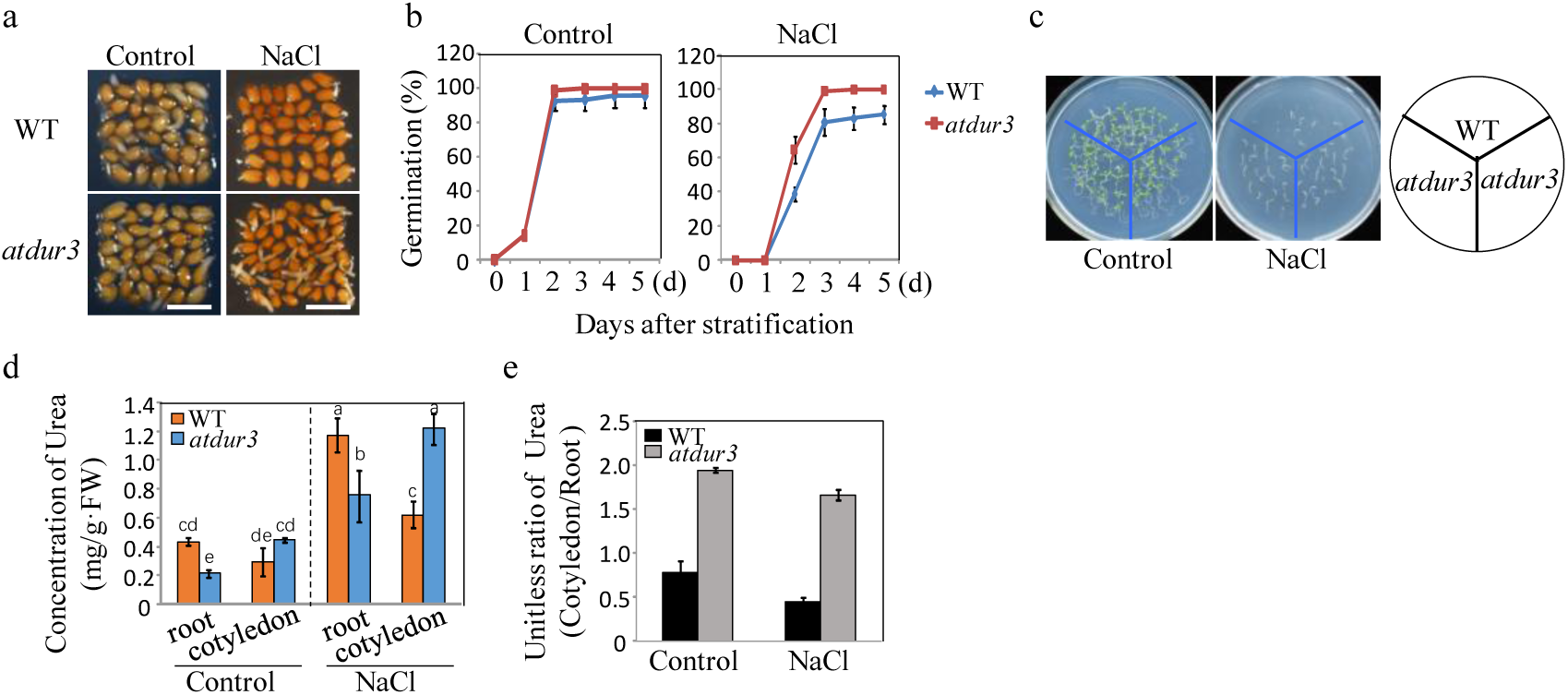
Arginine-derived urea transport and its role in salt-inhibited seed germination (SISG). (a) Seed germination phenotypes of WT and *atdur3* mutants were compared at 48 hours post-stratification at 4°C, focusing on radical emergence. Photographs captured under a stereomicroscope. Scale bar = 1 mm. (b) Germination rates of *atdur3* on ½ MS medium, containing 0 (control) or 135 mM NaCl, respectively, over time. (c) Growth comparison of WT and *atdur3* mutant seedlings on control and NaCl medium, photographed 10 days post-germination. (d) Urea content analysis in roots and cotyledons of 5-day-old seedlings grown on control or 135 mM NaCl medium. (e) Calculation of rrea (cotyledon/root) change ratio. In all experiments, seeds were freshly sowed and incubated under 16-h light and 8-h dark conditions at 22°C. The experiment was repeated three times; at least 30 seeds were counted in each replicate. The data was analyzed using a one-way ANOVA and Duncan’s post-hoc test, with different letters indicating significant differences (*P* < 0.05).

### Effects of arginine hydrolysis pathway on salt tolerance in other plants

Taken together, these results underscore the pivotal role of arginase-dependent arginine catabolic pathways in alleviating SISG in Arabidopsis. To investigate the conservation of this role across plants, we examined two major crops, *Oryza sativa* and *Glycine max*, along with two halophytes, *Chloris virgata* and *Puccinellia tenuiflora*. The seeds of *O.sativa* and *G.max* exhibited robust germination in the control medium with or without PPD. However, germination of seeds was notably inhibited in salt medium, a hindrance significantly improved by additional of PPD (Fig. 10a and b). Without PPD, germination rate was 20 ∼ 30% in the presence of NaCl, but with PPD supplementation, germination rates significantly increased to approximately 60 to 80% (Fig. 10a and b). Furthermore, root lengths of the seedlings were significantly shorter in NaCl medium compared to control conditions, yet supplementation with PPD in the NaCl medium led to increased root lengths (Fig. 10c and d). Similarly, studies on the typical halophytic plants demonstrated that under NaCl conditions, treatment with PPD significantly reduced salt-stress-induced inhibition of shoot and root growth compared to untreated counterparts (Fig. 10e). Notably, root lengths of samples treated with PPD under NaCl conditions were significantly higher than these without PPD treatment (Fig. 10f). These observations collectively suggest that blocking arginase-dependent arginine catabolism can enhance salt-tolerant seed germination in both crops and halophytes, underscoring the universal effect of arginine catabolism on plant salt tolerance.

**Figure 10.**
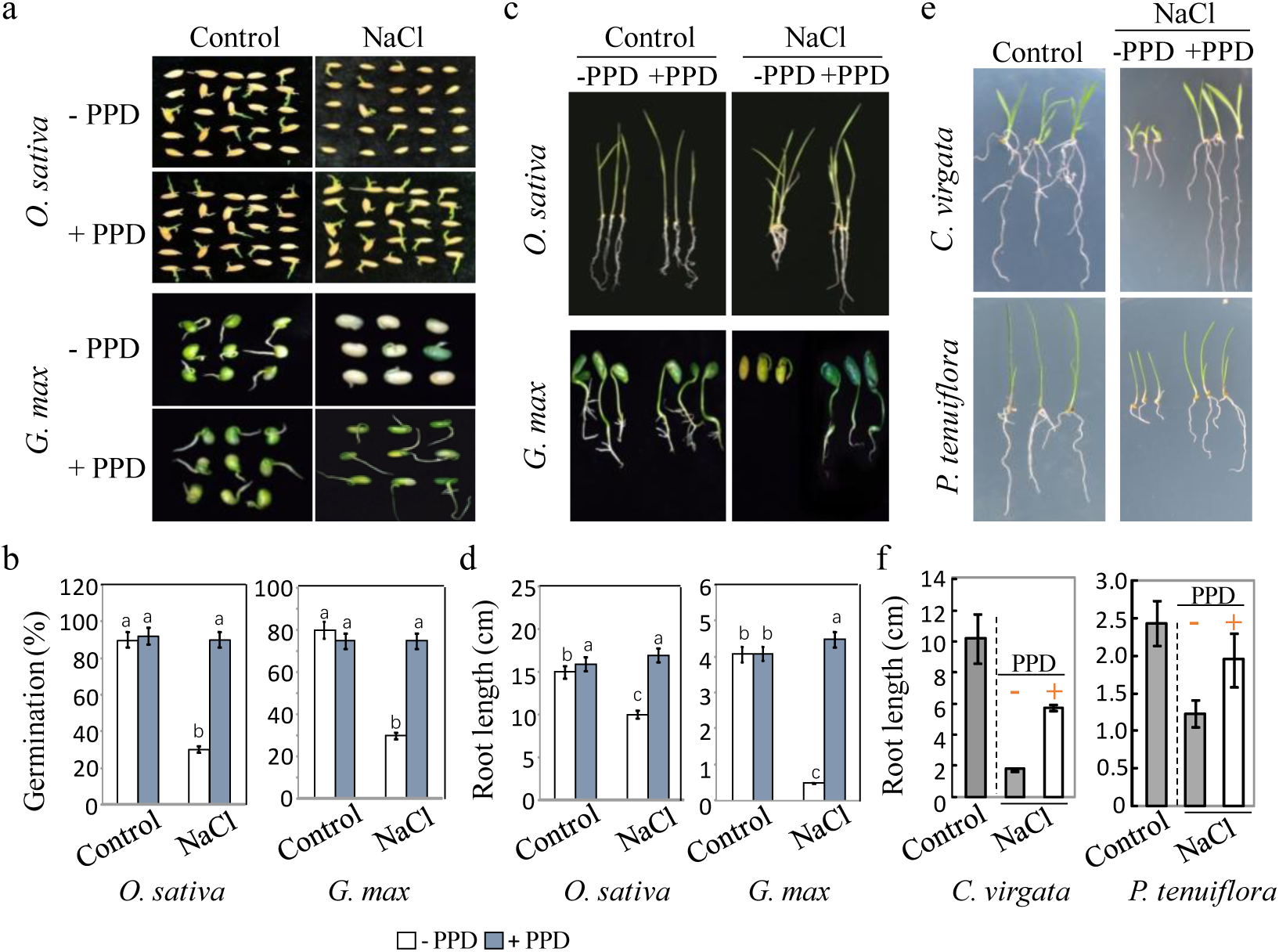
Effect of urea hydrolase inhibitor PPD, on the germination and seedling growth of *Oryza sativa* and *Glycine max*, *Chloris virgata* and *Puccinellia tenuiflora*. (a) Seeds of *O. sativa* and *G. max* were germinated on ½ MS medium (control; 0 mM NaCl) or ½ MS containing 150 mM NaCl with or without 15 µM PPD. Photographs were taken at 14 days after germination. (b) Germination rates of *O. sativa* and *G. max* plants in control or 150 mM NaCl with or without 15 µM PPD were calculated after 5 d sowing. (c) Seedling growth of *O. sativa* and *G. max* treated with and without PPD under salt stress, photographed 14 days after germination. (d) Root lengths was determined 14 d post-germination. (e) Seeds of *C. virgata* and *P. tenuiflora* were germinated 150 mM NaCl with or without 15 µM PPD, compared to control (0 mM NaCl). Representative morphological images of the treated seedlings 14 d post-stratification. (f) Root lengths was determined 14 d post-germination. The experiment was repeated three times with at least 30 plants per replicate. PPD, an inhibitor of urea hydrolase and a downstream metabolite of arginine.

## Discussion

The loss of food due to farmland salinization is staggering, with an estimated economic impact of exceeding $27 billion annually (Zörb et al., 2019; Kaleem et al., 2018). Failure of seeds to germinate stands as a significant contributor to crop loss, particularly under salt stress conditions, which not only delays germination but also reduces the overall percent of successful germination (Kojima et al., 2006; Kazachkova et al., 2016). Therefore, developing effective strategies to mitigate the detrimental effects of salinity on seed germination and seedling establishment is imperative. Here we focused on the effect of seed reserve nitrogen mobilization on salt-induced inhibition of seed germination (SISG). Arginase, a key enzymes in seed nitrogen storage mobilization, has garnered considerable attention in recent efforts aiming at understanding plant responses to abiotic stresses. Arginase-deficient mutants have been extensively utilized because of the competition among three enzymes, ARGAH, NOS and ADC, for a common substrate, arginine. It has been proposed that blocking arginase activity indirectly enhances the NOS and ADC pathways, thereby increasing the plant defense response mediated by NO and PA (She et al., 2017; Wang et al., 2011a; Flores et al., 2008). Since these explanations may be inadequate, here we clearly indicated that by employing experiments invovling arginase inhibitors, arginase gene mutants, as well as PPD and urease mutants, the arginine hydrolysis pathway (a two-step hydrolysis reaction) is directly involved in SISG (Fig. 1 and 2). This finding not only challenges existing explanations but also underscore the significance of seed reserve nitrogen mobilization in the context of SISG.

How exactly does the arginine hydrolysis pathway trigger SISG? Our hypothetical model, summarized in Fig. 8, elucidates the following process. The arginine hydrolysis pathway is implicated in triggering salt-induced inhibition of seed germination (SISG) through a series of interconnected physiological mechanisms. During seed germination, the activity of arginase and urease, especially particularly heightened under salt stress conditions, facilitates the conversion of of arginine to urea and then into the ammonia pool (Polacco et al., 2013; Siddappa et al., 2020; Kojima et al., 2006). Notably, arginase activity was significantly up-regulated under salt stress (Fig. 3e) (Polacco et al., 2013; Siddappa et al., 2020). This induction is further supported by strong evidence demonstrating the salt-induced expression of arginase and urease genes during seed germination (Polacco et al., 2013; Siddappa et al., 2020; Kazachkova et al., 2016; Lai et al., 2020), and triggers significant accumulation of urea and NH_4_ ^+^ (Fig. 5a and b). Together, these results indicate that the arginine hydrolysis pathway is induced by salt stress.

Interestingly, blocking the activity of arginase or urease necessarily triggers the accumulation of arginine or urea, respectively, yet paradoxically alleviates SISG. Thus, the toxic effects of arginine or urea accumulation can be excluded as a cause of SISG. Our results suggest that blocking arginase or urease activity leads to a decrease in NH_4_ ^+^ levels, which effectively mitigates SISG. This finding challenges our previous hypothesis that NH_4_ ^+^ accumulation due to urea hydrolysis under salt stress triggers SISG (Bu et al., 2015). However, our new results showed that the addition of ammonium to the medium under salt stress, on the contrary, effectively alleviated SISG (Fig. 5d and Supplementary Fig. 3). This new unexpected experimental results refutes this seemingly plausible hypothesis that ammonium accumulation (ammonia toxicity) resulting from excessive urea hydrolysis is sufficient to trigger SISG.

Recent studies have shed light on the role of glutamine synthetase 2 (GLN2), primarily located in plastid, in inducing acid stress through assimilation of NH_4_ ^+^, which exerts toxic effects on plants (Hachiya et al., 2021). In addition, our results showed that MSX treatment fails to alleviate SISG under salt stress, and interestingly, SISG shows a negatively correlation with GS activity (Fig. 6). This indicates that assimilation of NH_4_ ^+^ by GS may serve as an adaptive strategy for SISG. These results hint at the potential of cell acidification in alleviating SISG, which may explain why the addition of ammonium can effectively alleviate SISG. Traditionally, the hydrolysis of urea has been considered to produce ammonia and CO_2_; in fact, the urea hydrolysis in the two-step hydrolysis reaction of the arginine hydrolysis pathway, “(NH_2_)_2_CO + 3H_2_O → 2NH_4_ ^+^ + HCO_3_ ^−^ + OH^−^”, it is indeed an alkalinization reaction. In addition, further hydrolysis of bicarbonate will also produce hydroxide “HCO_3_ ^−^ + H O ⇆ H CO + OH^−^”, leading to further alkalization. Further investigation revealed that salt stress leads to an increase in cytoplasmic pH of seed radical cells, which results in SISG (Fig. 7). Overall, these hypothetical models suggest that blocking arginase or urease activity under salt-stressed conditions can effectively mitigate the events leading to SISG, both depending on whether the alkaline hydrolysis reaction of the core metabolite urea occurs or is enhanced, providing new insights into SISG (Fig. 7).

Salt inhibition of seed germination has been attributed to various signaling mediators and pathways (Zörb et al., 2019; Kaleem et al., 2018; Zhu et al., 2002; Zhu et al., 2016). Na^+^ toxicity is thought to be one of the main factors triggering SISG, as evidenced by the mitigation of SISG in the salt-sensitive mutant *sos3* by blocking the arginine hydrolysis pathway. This result implies that initiation of the arginine hydrolysis pathway exerts an additional effect on SISG (Fig. 4). Excessive hydrolysis of arginine leads to the accumulation of urea, and further hydrolysis of urea, coupled with elevated intracellular pH, leads to SISG events. Therefore, we posit that, in addition to Na^+^ toxicity, SISG is also instigated by elevated intracellular pH resulting from urea hydrolysis. salt sensitivity of *sos3* can be ameliorated by blocking urease activity and reducing intracellular pH under salt stress.

Urea is a plant metabolite derived from arginine hydrolysis or root uptake. In addition to the enzyme reaction required for urea hydrolysis, there are related urea transport steps in the mobilization process of nitrogen storage in seeds. In general, urea can be transported across plant cell membranes by high or low-affinity transporters (Polacco et al., 2013; Kojima et al., 2006; Bohner et al., 2015). On the cellular and mitochondrial membranes, low-affinity urea transporters are mediated by PIP/TIP or TIP aqueous proteins, respectively (Kojima et al., 2006; Bohner et al., 2015). The Dur3 mediates the high-affinity transport of urea. The Dur3 mRNA level increased significantly during seed germination (Bu et al., 2015; Van Zelm et al., 2020). According to our initial study, blocking the transport of free urea generated by the arginine hydrolysis pathway may also alleviate SISG (Fig. 9), and AtDur3 may be involved in long-distance urea transport from cotyledons to radicles. Urea transport and the arginine hydrolysis pathway may synergistically affect SISG.

In summary, our study elucidates the hydrolysis reaction of arginine-derived urea in mobilizing stored nitrogen in seeds and its role in triggering SISG by raising the pH levels in seed radicle cells. Since this evidence are primarily derived from Arabidopsis studies, the application of PPD treatment was tested on a diverse range of plants including glycophyte, halophyte, graminaceous crops, and legumes, which also followed the same underlying principles. This suggests that our observations may be applicable across the seed plant community (Fig. 10), indicating a broader relevance of our conclusions. Given that over half of the global nitrogen fertilizer is applied to crops in the form of urea, which adversely affects seed germination, the implications of urea metabolism and its transportation in relation to urea fertilizer use in agriculture warrant further exploration. Studies on the enzymes of arginine metabolism and the regulatory mechanisms that direct the allocation of arginine-derived nitrogen to defend against stress will enhance our understanding of the essential intermediates involved. Additionally, examining the developmental switches between nitrogen storage and remobilization could help to improve crop cultivation conditions and practices. Such studies will not only pivotal for optimizing crop cultivation conditions but also for mitigating the negative economic and environmental ramifications associated with excessive urea application in agriculture.

## Methods

### Plant materials and growth conditions

All experiments were performed with *Arabidopsis thaliana* Columbia (Col) wild-type plants and mutants in the Col background. The T-DNA insertion line SALK_057987 (*atargah1*) was ordered from the ABRC (Ohio State University), and homozygous lines were confirmed by genome PCR using primers 057987-LP/RP or 036318-LP/RP and a T-DNA primer LBP (Supplementary Table 1). Other lines, including SALK_038002 (*aturease*), SAIL118_C11 (*atargah2*), and *sos3*, have been previously described (Sessions et al., 2002; Zhu et al., 1998). The double mutants *sos3*/*aturease* was produced by crossing, whereas *atargah1/atargah2* was produced using CRISPR-Cas9. Homozygous plants were selected from F_2_ populations, and F_3_ plants or further generations were used for analysis.

Wild-type and mutant plants were cultivated concurrently, with seeds collected simultaneously. Seeds were sown on half-strength MS (½ MS) medium containing 0.8% (w/v) agar and plates were kept in the dark at 4°C for 2 days to break dormancy (stratification). Subsequently, they were transferred to a climate chamber for 14-day cultivation period (22°C; 16/8 h light/dark; light intensity of 40 mmol m ^-2^ s ^-1^). Seedlings were then moved to soil and cultured in a greenhouse under a photoperiodic cycle of 16 h light and 8 h dark photoperiod at 22°C. Fully developed and ripened brown siliques were collected for analysis.

### Seed germination assay

Seeds stored for a period ranging from 2 weeks to 3 months at room temperature were used for germination test. The after-ripened seeds were sterilized using 75% (v/v) ethanol and 10% (v/v) NaClO for 1 min, followed by three times washes in sterile water. Subsequently, they were plated on sterile filter paper to air dry. Following sterilization, the seeds were sown on solid medium consisting of ½ MS supplemented with varying concentrations of NaCl (0, 135 mM), *^N^*G-Hydroxy-L-arginine (NOHA; Sigma, Germany; 0 – 5 µM), Phenyl phosphorodiamidate (PPD; Macklin, China; 0–7.5 µM), L-methionine sulfoximine (MSX; Sigma, Germany; 0, 3 µM). Salt concentrations were set at 0 mM for controls and 135 mM for treatments unless noted otherwise. Plates were stratified at 4°C in darkness for 2 days, then transferred to an illumination incubator at 22°C under 16 h light/8 h dark cycle for subsequent analysis. At least 30 seeds for each genotype were used in three biological replicates. The germination event was defined as the initial emergency of the radicle, observed and recorded at 48 h of incubation. The assessment of germination, cotyledon-greening test and radical growth used in the current study were as previously described (Bu et al., 2015). Seed germination rates were assessed daily in triplicates, with each plate containing over 35 seeds.

Seeds of wild-type *Oryza sativa*, *Glycine max*, *Chloris virgata* and *Puccinellia tenuiflora* were surface-sterilized in 1% NaClO solution for 10 min, followed by washing three times in sterilized distilled water. Then, 30 seeds were sown on ½ MS (i.e., half the concentration of regular MS) supplemented with NaCl (0 or 150 mM) and Phenyl phosphorodiamidate (PPD; 0 or 15 µM), then subjected to stress conditions at 22°C for 14 days. Germination was defined by an obvious emergence of the radicle through the seed coat, with counts taken 3 to 5 days post-sowing. The germination rate was calculated as follows: Germination rate (%) = (number of germinated seeds/total number of seeds) × 100%. Root length and fresh weight were measured after 14 days of cultivation. Data were analyzed using three biological replications, and statistical significance was determined using Duncan’s test.

### Constructs and plant transformation

CRISPR-Cas9 technology was employed to design specific sgRNAs targeting *ArabidopsisAtArgAH1* and *AtArgAH2*. The gRNA-U6 fragment, once amplified, was cloned into the M2CRISPR vector (14,847 bp). The resulting construct, named CR-PCR-*AtArgAH1AtArgAH2*, was then introduced into *Arabidopsis* Columbia wild-type plants using stable transformation with *Agrobacterium tumefaciens*. Genomic DNA was extracted from young leaves of transformed plants and amplified by PCR using primers flanking the target sites to confirm the introduction of mutations. The PCR products were sequenced to identify double mutants, *atargah1*/*atargah2*. The primer sequences were using primers *AtArgAH1*-FW/RV and *AtArgAH2*-FW/RV (Supplementary Table 1).

### Measurement of Arginase activity

To detect arginase activity, WT, mutant lines of Arabidopsis (*atargah1*, *atargah2*, and the double mutant *atargah1*/*atargah*, with deletions in *AtArgAH1*or *AtArgAH2*) were germinated for 3 days on media containing 0 or 135 mM NaCl, following a stratification period of 48 hours at 4°C in darkness. The arginase activity was measured using a commerical assay kit (Bioassay, USA) following the instructions supplied by the manufacturer. Briefly, approximately 0.1 g of tissue was homogenized in 1 mL of extraction buffer on ice. After centrifugation at 8000 g for 10 min, the supernatant was collected and placed on ice. For the assay, 10 µL of substrate buffer was added to the samples, and 200 µL urea reagent was added to the control tubes, respectively. After through mix, the samples were incubated at 25°C for 60 minutes. Arginase activity was quantified by measuring the absorbance of the supernatant at 430 nm.

### Extraction and quantification of urea

For urea extraction, 1 mL of 10 mM ice-cold formic acid with the addition of the urease activity inhibitor (PPD) were added to about 100 mg sample to avoid urea hydrolysis. Urea concentration was then quantified using a commercial assay kit (BioAssay Systems, USA) following the manufacturer’s instructions. Each sample mixture was vigorously vortexed twice before centrifuged at 16,000 rpm for 15 min at 4°C. The supernatant was carefully transferred to a fresh tube for analysis. Next, 200 µL of the provided color development reagent was added to each sample in a microcentrifuge tube. Tubes were incubated 20 min at room temperature, and optical density of each sample was measured at 520 nm to quantify urea concentration.

### Ammonium concentration assay

Cotyledons and roots of Arabidopsis plants were gently washed with ice-cold Milli-Q water, dried with tissue paper, and immediately frozen in liquid nitrogen. For extraction, 0.5 gram of frozen tissue was combined with 5 mL ice-cold extraction medium, and a small amount of quartz sand in a chilled mortar. The sample was then ground to a fine powder with a chilled pestle. Determination of the concentration of ammonium in plant tissue was conducted as previously described (Bu et al., 2015). NH_4_ ^+^ concentration was determined colorimetrically at 640 nm.

### Confocal Laser Scanning Microscopy

Arabidopsis seeds engineered to express PRpHluorinwere were germinated on solid MSat at 22°C for 3 days to facilitate pH measurement studies. The PRpHluorin gene was amplified using the specific primers (PRpHluorin-F and PRpHluorin-R) and then cloned into pBI121 vectors, which were modified to include the UBQ10 promoter. These constructs were introduced into WT Columbia-0 (Col-0) Arabidopsis plants via Agrobacterium-mediated transformation, employing the floral dip method followed by selection on kanamycin. For pH measurement in seedlings, fluorescent images were captured using a Zeiss LSM 880 confocal laser scanning microscope (CLSM) equipped with a 20x or 63x objective. The imaging was conducted in a sequential line scanning mode with settings described previously (Shen et al., 2013). Briefly, the emission (500 - 550nm) of pHluorin, triggered by sequential excitation with 488- and 405-nm lasers, was used to calculate the pH using the calibration curve. In vivo calibration was performed on a subset of the seedlings at each experiment’s conclusion, where they were immersed pH equilibration buffers for 15 minutes (Krebs et al.,2010). The buffers contained 50mM Mes-BTP (pH 5.2–6.4) or 50 mM Hepes-BTP (pH 6.8–7.6), supplemented with 50 mM ammonium acetate. The emission ratio was plotted against pH, and sigmoidal curves were fit to the data using a Boltzmann equation, as described previously (Gao et al., 2004; Schulte et al., 2006). For post-treatment pH calculation, 10 seedlings with 100 cells per root were analyzed. The selected area was represented by the yellow dotted boxes (Supplementary Fig. 5), and pH values were deduced from the in vitro calibration curves using ImageJ software. Statistical analyses were conducted using one-way ANOVA in Graphpad Prism. To visually represent the pH profile, the grayscale ratio images were converted to pseudocolored images using the ImageJ.

### Statistical analysis

For physiological and biochemical data, an analysis of variance was performed to investigate whether there was a significant difference between the samples. If a significant difference was found, a Duncan’s significant difference test was performed to determine the specific samples with significant differences.

## Acknowledgements

This work was supported by the Heilongjiang Province Government Postdoctoral Science Foundation (LBH-Q18008) awarded to Yuanyuan Bu. Further supported by the Program for Changjiang Scholars and Innovative Research Team in University (No. IRT17R99) awarded to Shenkui Liu. The funders had no role in study design. We are grateful to Professor Jinbo Shen (Zhejiang Agriculture and Forestry University, Lin’an, China) for providing PRpHluorin seeds and technical support for cytoplasmic pH measurement.

## Author Contributions

YY. Bu and SK. Liu designed the study. XY. Dong, RR. Zhang, XL. Shen, Y. Liu, S. Wang and YY. Bu performed the experiments and analyzed the data. YY. Bu, SK. Liu and T. Takano supervised the study and critically reviewed the manuscript. All authors read and approved the final manuscript.

## Abbreviations list

1/2 MS: One-half-strength Murashige and Skoog
WT: Wild type
SISG: Salt-induced inhibition of seed germination
GS/GLN: glutamine synthetase
NOS: nitric-oxide synthase
ADC: Arginine decarboxylase
ARGAH: arginine amidinohydrolyase
NO: Nitric Oxide
PA: polyamine
PPD: phenyl phosphorodiamidate
NOHA: *N*^G^-hydroxy-L-arginine
MSX: L-methionine sulfoximine
GOGAT: Glutamate synthasis
SOS: Salt Overly Sensitive

